# vmPFC drives hippocampal processing during autobiographical memory recall regardless of remoteness

**DOI:** 10.1101/2020.04.27.063875

**Authors:** Cornelia McCormick, Daniel N. Barry, Amirhossein Jafarian, Gareth R. Barnes, Eleanor A. Maguire

## Abstract

Our ability to recall past experiences, autobiographical memories (AMs), is crucial to cognition, endowing us with a sense of self and underwriting our capacity for autonomy. Traditional views assume that the hippocampus orchestrates event recall, whereas recent accounts propose that the ventromedial prefrontal cortex (vmPFC) instigates and coordinates hippocampal-dependent processes. Here we sought to characterise the dynamic interplay between hippocampus and vmPFC during AM recall to adjudicate between these perspectives. Leveraging the high temporal resolution of magnetoencephalography, we found that the left hippocampus and the vmPFC showed the greatest power changes during AM retrieval. Moreover, responses in the vmPFC preceded activity in the hippocampus during initiation of AM recall, except during retrieval of the most recent AMs. The vmPFC drove hippocampal activity during recall initiation and also as AMs unfolded over subsequent seconds, and this effect was evident regardless of AM age. These results re-cast the positions of the hippocampus and the vmPFC in the AM retrieval hierarchy, with implications for theoretical accounts of memory processing and systems-level consolidation.

## Introduction

Our past experiences are captured in autobiographical memories (AMs). Functional MRI (fMRI) studies over many years have shown that the hippocampus and ventromedial prefrontal cortex (vmPFC) are among a distributed set of brain areas that is consistently engaged during the retrieval of such memories (reviewed in Maguire 2001; Svoboda et al. 2006; McDermott et al. 2009; Spreng et al. 2009). In addition, decoding approaches have revealed that patterns of fMRI activity associated with specific AMs are evident in the hippocampus, irrespective of the age of a memory (Bonnici et al. 2012; Bonnici and Maguire 2018), while detectability of individual memories increases in vmPFC as they become more remote (Bonnici et al. 2012; Barry et al. 2018; Bonnici and Maguire 2018).

Neuropsychological studies complement the fMRI findings, showing that patients with bilateral hippocampal damage are either unable to recall AMs at all (Scoville and Milner 1957; reviewed in Clark and Maguire 2016; McCormick et al. 2018a) or retrieval is significantly impoverished (Viskontas et al. 2000; Addis et al. 2007; St-Laurent et al. 2011; St-Laurent et al. 2014). vmPFC lesions are also associated with significant impairment of AM recollection (Della Sala et al. 1993; Bertossi et al. 2016; reviewed in McCormick et al. 2018a), and in addition can provoke confabulation. This involves the production of false AMs that patients believe to be true, perhaps due to an inability to select the appropriate components of memories and inhibit those that are irrelevant (Moscovitch and Melo 1997; Ciaramelli et al. 2006; Gilboa and Marlatte 2017).

While fMRI co-activation studies and neuropsychological findings present clear evidence that the hippocampus and vmPFC are involved in supporting AM recall, they do not inform about whether the two regions actually interact in the service of retrieval. Structural MRI studies have shown that the hippocampus and vmPFC are connected (Catani et al. 2012; Catani et al. 2013), and individual differences in the microstructure of a key fibre connection, the pre-commissural fornix, correlate with the richness of autobiographical memories (Williams et al. 2020). fMRI functional connectivity analyses have gone a step further and documented correlated activity between the hippocampus and vmPFC during the retrieval of AMs (e.g. Addis et al. 2004; McCormick et al. 2015; Robin et al. 2015; Inman et al. 2018; McCormick et al. 2018b; Sheldon and Levine 2018). A small number of fMRI studies have also examined effective connectivity in the context of AM retrieval, by investigating whether one brain region exerted influence over the other, allowing for a deeper understanding of the causal dynamics that enable AM recall. For example, using dynamic causal modelling (DCM; Friston et al. 2003), St Jacques et al. (2011) reported that medial prefrontal cortex drove activity within the distributed AM recall network, particularly during the elaboration phase in the seconds following initial recall. Nawa and Ando (2019) also found that the vmPFC influenced the hippocampus during AM recall elaboration. In both studies non-specific generic cues were used to elicit memories. It has been proposed that vmPFC may lead retrieval particularly in circumstances where cues are generic and lack specificity (Robin and Moscovitch 2017), which could account for these findings.

While fMRI studies such as these have increased our understanding of how AM retrieval is supported by the brain, fMRI has some fundamental constraints. It is not a direct measure of neural activity and is therefore slow – the haemodynamic response is in the order of ∼6 seconds. Consequently, it cannot inform about the millisecond neural dynamics that are key to elucidating the mechanisms underpinning AM recall. It also precludes examination of the earliest stage of AM retrieval, which is critical for the success of the ensuing recall process. By contrast, magnetoencephalography (MEG) provides a direct measure of neural activity with millisecond temporal resolution. Measuring neural dynamics from deep sources such as the hippocampus was initially thought to be beyond the sensitivity of MEG. However, over the years, progress in MEG modelling and validation though simulation, invasive recordings and fMRI, has provided ample evidence that deep brain sources can be measured using MEG (e.g. Poch et al. 2011; Kaplan et al. 2012; Staudigl and Hanslmayr 2013; Dalal et al. 2013; Backus et al. 2016; Meyer et al. 2017).

To the best of our knowledge, just two AM retrieval MEG data sets have been reported. Fuentemilla et al. (2014; the same data were also reported in Fuentemilla et al. 2018) had eight particpants recall AMs that were between two and seven months old. Hebscher et al. (2019; the same data were also reported in Hebscher et al. 2020) used continuous theta burst stimulation of the precuneus during the recall of memories that were under one year old. Both data sets showed phase synchronisation between the medial temporal lobe and the medial prefrontal cortex during AM retrieval, providing further evidence that these two areas seem to cooperate in facilitating recall of the past.

Despite the potential of MEG for elucidating AM-related hippocampal-vmPFC interactions, numerous questions remain unanswered. Key among them is whether the hippocampus or vmPFC engages first during the very earliest initiation phase of the AM retrieval process, and whether one region drives activity in the other. This, currently missing, information is essential for building a mechanistic understanding of how neural responses give rise to the recall of past events. Similarly, after initiation of AM recall, how do hippocampus and vmPFC influence each other to facilitate recall of the unfolding AM over subsequent seconds? The fMRI DCM findings (St Jacques et al. 2011; Nawa and Ando (2019) suggest that vmPFC might drive hippocampus during this elaboration stage, but would this still be the case if the memory cues were highly specific? Also critical to consider is whether, and how, AM remoteness affects hippocampal-vmPFC neural dynamics.

Previous studies have not considered any of these issues. In the current study, therefore, we addressed these questions by using MEG to interrogate neural activity in the hippocampus and vmPFC in healthy adults as they vividly recalled AMs triggered by specific cues, where the age of memories was also carefully manipulated.

Considering our hypotheses, different views exist about the roles of the hippocampus and vmPFC and their interactions in supporting AMs. For instance, some accounts place the hippocampus at the heart of AM retrieval and believe it recruits neocortical regions in the service of this endeavour (Teyler and DiScenna 1986; Teyler and Rudy 2007). If this is the case, then the hippocampus should engage first during initial AM recall, and drive activity in vmPFC during this phase, and perhaps also during the subseqent seconds of recollection.

Another perspective proposes that the frontal cortex may guide memory search which may then enable hippocampally-mediated recovery of a memory (Moscovitch 1992; Shallice and Burgess 1996; Moscovitch and Melo 1997), particularly in the context of non-specific cues (Robin and Moscovitch 2017). In another account, the vmPFC and not the hippocampus is held to initiate imagery-rich mental events such as those involved in AM retrieval even when cues are specific (McCormick et al. 2018a; Barry and Maguire 2019a,b). The occurrence of confabulation, along with reduced instigation of spontaneous thoughts more generally following vmPFC damage (Kleist et al. 1940; Bertossi and Ciaramelli 2016; McCormick et al. 2018a), accords with the vmPFC being involved at the earliest point of AM recall initiation. Moreover, a recent MEG study involving the imagination of visual scenes, which Ams typically comprise, documented engagement of the hippocampus and vmPFC and found that the latter initiated the construction of these mental scenes and drove activity in the hippocampus (Barry et al. 2019a). Within this account, therefore, the vmPFC should engage first during initial AM recall in response to specific cues, and drive activity in the hippocampus during this phase, and perhaps also during the subsequent seconds of recollection. Whilst we favour this view, our paradigm enabled adjudication between the different perspectives, given the clearly contrasting predicted outcomes.

There is also debate about the nature of hippocampus and vmPFC involvement in retrieval as a function of AM age. The hippocampus in particular is regarded as being required solely for the retrieval of recent AMs (Squire 1992), or for recalling detailed and vivid AMs in perpetuity because there is a permanent trace stored in the hippocampus (Nadel and Moscovitch 1997; Robin and Moscovitch 2017; Moscovitch and Nadel 2019), or for recollecting vivid and rich AMs of any age because it constructs the scenes that are central to re-experiencing past events (Maguire and Mullally 2013; Zeidman and Maguire 2016; Barry and Maguire 2019a,b).

Barry and Maguire (2019a,b; but see Moscovitch and Nadel 2019) recently reviewed data from the cellular to the systems level and found little evidence that the hippocampus stores anything in the longer-term, but nevertheless seems critical for recalling AMs even when they are very remote. They concluded that the most parsimonious explanation for this apparent paradox involves the hippocampus encoding autobiographical experiences and storing AMs in the short-term. However, its role in their subsequent retrieval then becomes one of reconstructing the scenes that comprise these unfolding events, with orchestration of this process likely coordinated by neocortical regions such as the vmPFC. On this basis, the prediction would be that during recall of recent AMs (defined as less than one month old; Bonnici et al. 2012; Barry et al. 2018), there will be no evidence of vmPFC activity preceding that of the hippocampus because complete representations of AMs will still be available in the hippocampus. In contrast, for AMs that are not recent, a lag in engagement between vmPFC and hippocampus will be apparent, with the former leading. Here, we elucidate the neural dynamics associated with AM remoteness using MEG to provide new leverage on key debates involving theories of memory consolidation.

## Materials and Methods

### Participants

Eighteen right-handed healthy adults (10 males, mean age 31.6 years, SD 5.0) with normal vision and hearing participated in the experiment, which involved one AM selection meeting and one MEG scan 14 days later. All participants gave written informed consent. The study was approved by the University College London Research Ethics Committee.

### Autobiographical Memory Selection

At the outset, the experimenter explained to the participants the type of autobiographical memories we sought to include, namely, a memory had to be specific in time and place, a unique event, and vividly recollected. Several examples were provided by the experimenter, such as a specific event during a vacation or the wedding of a friend. Participants were asked to select 50 autobiographical memories that met these criteria. Motivated by Barry et al. (2018), they were asked to retrieve 12 AMs that were less than one month old (<1M), 12 that were between four and twelve months old (4-12M), 12 that were between sixteen and twenty months old (16-20M), and 12 AMs that were between two and five years old (2-5Y). Two additional events were selected for practice purposes. Participants were instructed to choose memories that they recalled very clearly. They were also told that they would need to describe the memory to the experimenter, so they should not include very private memories. In addition, across the 50 AMs, they were asked to recollect memories involving different people, different places, and different topics, to minimise interference between events. Memories were allowed to vary in terms of their personal significance.

Participants were provided with a sheet of paper on which they could make notes about the AMs and the age of the memories in their own time. When they had finished (about one hour later), all memories were reviewed together with the experimenter to check that they met the criteria for unique AMs. For each memory, the participant was instructed to give a short account of that event to ensure that it could be recalled in rich detail. Participants were then asked to rate, on a 5-point scale, a memory’s vividness (only vivid and very vivid answers were accepted), how easy it was to recall (only easy and very easy answers were accepted), its personal significance (all answers were accepted), emotional valence and intensity (only neutral and slightly positive answers were accepted), rehearsal frequency (only never, very little and occasionally answers were accepted), whether it was had first or third person perspective (only first person events were accepted), and active or static (only active events were accepted). The vast majority of the memories offered by participants were suitable for inclusion in the experiment; very few were not. If a memory did not meet the criteria, the participant was asked to provide a different memory for that age bracket. Of note, during a pilot phase of the study we collected both vividness and level of detail subjective ratings. However, participants did not seem to differentiate between the two which led to a precise mirroring of the results. We, therefore, opted to include just one of these ratings in the study proper. We chose vividness given the good evidence that hippocampal activity correlates with the vividness of autobiographical memory retrieval (e.g. Sheldon and Levine 2013; see also Clark and Maguire 2020). Each autobiographical memory was given a unique two-word title by the participant that was subsequently recorded and played to the participant during the MEG scan. No titles started with the same word (i.e. “Chris’ wedding” and “Chris’ birthday”), to prevent confusion. This AM selection meeting lasted between two to three hours, including breaks.

### Task and Procedure

After the AM selection meeting, the 50 two-word titles for each participant were recorded and cut into short audio clips using Audacity (https://sourceforge.net/projects/audacity). Upon arrival for the MEG scan two weeks after the selection meeting, participants were allowed to study the 50 two-word titles one more time, so they knew which events they had to recall in the scanner. Once positioned in the MEG scanner, the task was explained to them (Figure 1). There were four practice trials followed by the experiment proper. In the scanner, each trial started with a 1 sec visual cue “Please close your eyes!” presented on a screen. Participants closed their eyes immediately and waited for an auditory cue which followed a jittered duration (mean 3 sec, SD 1.0). This cue was either one of the titles that prompted participants to recall an AM, or a two-word instruction related to a baseline condition which involved counting (e.g., “Count 3s”). Participants were asked to retrieve the AM in as vivid and detailed a manner as possible or, for the counting condition, to count silently in steps of 3’s, 7’s, or 9’s, until a beep was sounded after 10 sec, which instructed participants to open their eyes. They were then presented with a screen prompting them to rate, via a key pad and within 3 sec, how successful they were at engaging in the task during that trial, for example, whether they successfully re-experienced the AM for the entire trial time (response 1), whether they started to recall the AM for part of the trial time but then got distracted (response 2), or whether they could not recall the AM at all either because they did not understand the cue, did not remember the AM, or were otherwise distracted (response 3). The stimuli were delivered aurally via MEG-compatible earbuds using the Cogent toolbox (www.vislab.ucl.ac.uk/cogent.php) running in Matlab (version 2012b).

**Figure 1.**
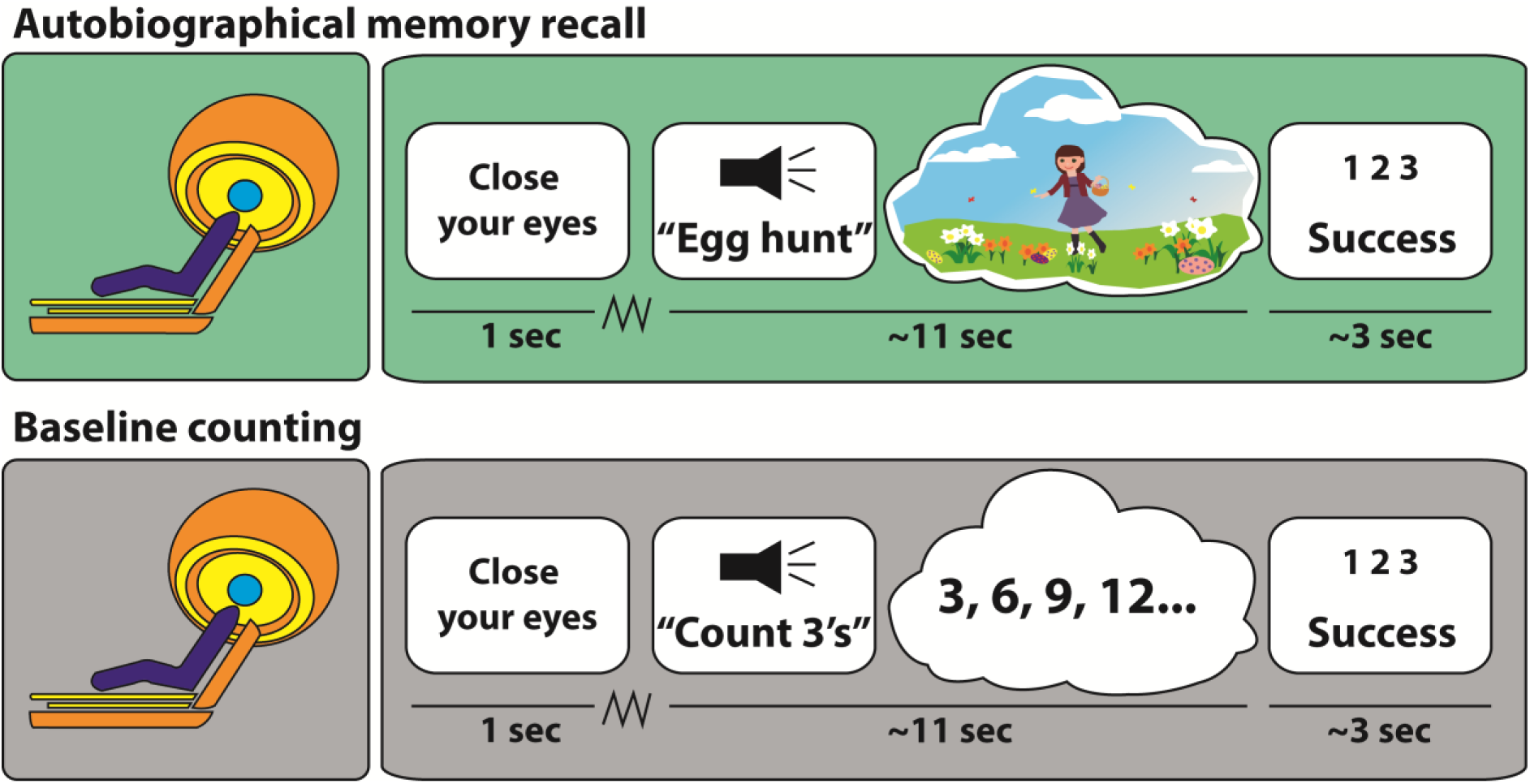
Example trials. Participants saw a cue instructing them to close their eyes and they then waited for an auditory cue which followed a jittered duration (jagged line). Upper panel: they heard a two-word cue relating to a specific AM – this example concerned an Easter egg hunt. They then had 10 sec to recall the AM in as vivid and detailed a manner as possible. At the end of each trial, participants heard a beep alerting them to open their eyes and rate the success of the retrieval (1=engaged fully for the entire trial; 2=partly engaged during the trial; 3=did not engage at all during the trial). Lower panel: the counting baseline trials had the same timing, but instead of recollecting an AM, participants had to mentally count in steps of either 3’s, 7’s or 9’s.

### MEG Data Acquisition and Preprocessing

A CTF Omega whole-head MEG system with 273 second order gradiometers recorded data at a sampling rate of 1200 Hz. MEG data were epoched into 5 sec AM retrieval and counting periods, baseline corrected, and concatenated across sessions.

### Behavioural Data Analysis

Comparison across AMs of different ages for each of the five ratings was performed using a repeated measures one-way ANOVA and deemed significant when p<0.05. Where an ANOVA was significant, Sidak correction post-hoc t-tests were used to account for multiple comparisons. We also applied Bonferroni correction across the five repeated measures ANOVAs, with the results deemed significant at p<0.01. Graphpad Prism version 6 was used for the statistical analyses.

### MEG Data Analysis – Source Reconstruction

All MEG analyses were performed using SPM12 (www.fil.ion.ucl.ac.uk/spm). Source reconstruction was performed using the SPM DAiSS toolbox (https://github.com/spm/DAiSS). A Linearly Constrained Minimum Variance (LCMV) beamformer was used to estimate differences in power between the two conditions, AM retrieval and the counting baseline. This filter uses a series of weights to linearly map MEG sensor data into source space to estimate power at a particular location, while attenuating activity from other sources. For each participant, a single set of filter weights was constructed based on the data from the two conditions within a broadband signal (1-30 Hz) and a 0 to 5000 msec peri-stimulus window. Analysis was performed in MNI space using a 5 mm grid, and coregistration was based on nasion, left and right periauricular fiducials. Coregistration and the forward model were computed using a single-shell head model (Nolte 2003). Power was estimated with one image per condition being generated for each participant. These images were entered into a second-level random effects paired t-test in SPM to investigate power differences between conditions. Images were thresholded at p<0.001 uncorrected (given our strong a priori hypotheses about vmPFC and hippocampus) and a cluster extent of >50 consecutive voxels.

For subsequent analyses, based on the source reconstruction results and our a priori specific interest in the hippocampus and vmPFC, time series data of broadband (1-30 Hz) activity during the 5000 msec of AM retrieval and the baseline counting condition were extracted as an average over voxels from anatomical regions of interest (ROIs) for the whole left hippocampus and the vmPFC using the LCMV beamforming algorithm. The masks for the ROIs were created using FSL v6.0 (https://fsl.fmrib.ox.ac.uk/fsl).

### Event-Related Analysis

Event related analysis followed the standard procedure. First, MEG data were epoched based on a peri-stimulus time window of −500 to 1000 msec. Using a Butterworth filter, data were bandpass filtered between 1 and 30 Hz. Data were then downsampled to 200 Hz and finally averaged using the robust averaging algorithm implemented in SPM12. Resulting files were exported from Matlab to Graphpad Prism for illustration purposes. In order to analyse the temporal order of vmPFC and hippocampal activity, maximum responses and their respective time positions were extracted for each condition (i.e., AM retrieval, baseline counting, and the different AM age categories) for each participant. Statistical analyses of these values were performed with Student’s paired t-test with Bonferroni correction performed across the four t-tests, with the results deemed significant at p<0.01. The latencies of responses across vmPFC and hippocampus were also examined using a repeated measures one-way ANOVA using Sidak correction post-hoc t-tests to account for multiple comparisons.

### Dynamic Causal Modelling (DCM) of Electrophysiological Data

DCM is a method used to specify, fit and compare biophysically-informed models based upon features (e.g., event-related signals) of neuroimaging data. The generative model of electrophysiological recordings in DCM is based upon interconnected neuronal mass models (NMM) each of which is specified via interactions between neuronal populations (e.g., excitatory, inhibitory and pyramidal cells). Each neuronal population within an NMM converts receiving intrinsic (within a region) and/or extrinsic (from a distal region) weighted (i.e., effective connectivity) firing rate inputs (sigmoid transformation of membrane potentials) to a postsynaptic potential (through convolution operator as a model of synaptic transmission) which then represents the input to other populations (Moran et al. 2013). Here we used a convolution-based NMM to form hypotheses regarding differences in (extrinsic) effective connectivity (Moran et al. 2013). The extrinsic connectivity in our model was characterised as forward or “bottom up” if projections were to the middle layers of the cortex, backward or “top down” if projections targeted deep and superficial layers, or lateral if projections innervated all layers (Felleman and Van Essen 1991). We could, therefore, test biologically plausible models based on known structural connections between our two ROIs that differed in terms of which connections were functionally modulated by the experimental task.

The DCM pipeline begins by fitting biologically-informed models to the features of the neuroimaging data through optimisation of model evidence (which accounts for a balance between model accuracy and complexity). The ensuing estimation using DCM includes posterior estimate of parameters and model evidence (also known as free energy) which in turn are used for model comparison across participants.

To assess which model best explained the observed data at a group level, random effects Bayesian model comparison (Stephan et al. 2009) was performed which compares the evidence for each model across participants, and generates the probability of one model being the winning model. To assess the quality and consistency of the model fit, we generated the log Bayes factor for each participant separately by computing the difference between the log evidence of the two models. For the analysis of different memory ages, we extracted the value of free energy for each participant as well as the connectivity strength parameter for each contrast for each participant. This technique allowed us to examine whether connectivity strength differed across the four categories of AM age. These data were then analysed using a repeated measures one-way ANOVA and deemed significant when p<0.05, with Sidak correction post-hoc t-tests employed to account for multiple comparisons.

#### DCM for Event-Related Signals

DCM of event-related signals (Garrido et al. 2007) was used to infer the effective connectivity between hippocampus and vmPFC during the initiation of AM retrieval. DCM for event-related signals maximises data fit between two or more predefined event-related signals. Two simple models were specified, namely, one where vmPFC influenced hippocampal activity and the other where the hippocampus influenced vmPFC activity. We modelled data in the period between 50 and 300msec after stimulus onset in order to capture the very early phase of AM retrieval initiation. After DCM for event-related signals maximised the fit of the neural data to the two models, random effects Bayesian model comparison for group studies was used to determine the winning model.

#### DCM for Cross Spectral Densities (CSD)

DCM for CSD (Kiebel et al. 2009; Moran et al. 2009) was used to infer effective connectivity between the vmPFC and hippocampus during AM retrieval and the counting baseline based on spectral features of the MEG data during the first 5 sec after cue onset. DCM for CSD involves specifying the direction of inter-regional information flow and fitting this model (formally, our biological hypothesis) to the spectral contents of MEG data. Multiple possible models are fitted to the data and the ensuing inferred models are then compared to ascertain the best explanation for the experimental observations. Here, we specified two simple models, one where the vmPFC influenced hippocampal activity and the other where the hippocampus influenced vmPFC activity, and we then compared their evidence over the participant group.

## Results

### Characteristics of the AMs

The AMs included in the experiment were selected because they were recounted in rich detail, and met the criteria required by the experiment. This meant that the memories were judged to be very vivid (mean=4.4, SD=0.3, scale 1-5, where 5 was the highest vividness score), easy to recall (mean=1.1, SD=0.2, scale 1-5, where 1 was the easiest recall score), of average personal significance (mean=2.9, SD=0.6, scale 1-5, where 5 was the most personally significant score), of neutral to slightly positive emotional valence (mean=3.7, SD=0.3, scale 1-5, where 5 was the highest positive valence score), and not frequently rehearsed (mean=2.5, SD=0.6, scale 1-5, where 5 was the most often rehearsed score). During MEG scanning, which happened two weeks after the AM selection meeting, AM retrieval was rated as successful (mean=1.1, SD=0.1) and no AM retrieval trial was rated as unsuccessful.

### vmPFC and Hippocampal Neural Dynamics Support AM Retrieval

We first determined which brain regions were engaged during AM retrieval by estimating the difference in broadband power (1-30 Hz) between the AM and counting baseline conditions in source space. Significant power changes were observed in just two brain regions, which coincided with our areas of interest – the vmPFC (peak MNI coordinate: 14, 60, −10, t=3.51) and left anterior hippocampus (peak MNI coordinate: −20, −6, −24, t=3.42; Figure 2). Changes in both regions represented an attenuation of power during AM retrieval. The direction of this power change aligns with numerous previous reports of decreased power during other types of memory tasks and the generation of scene-based mental imagery (Fellner et al. 2017; Barry et al. 2019a). The opposite contrast revealed attenuation of power in the right superior temporal cortex (peak MNI coordinate: 60, −30, 6, t=4.07) during the counting baseline condition.

**Figure 2.**
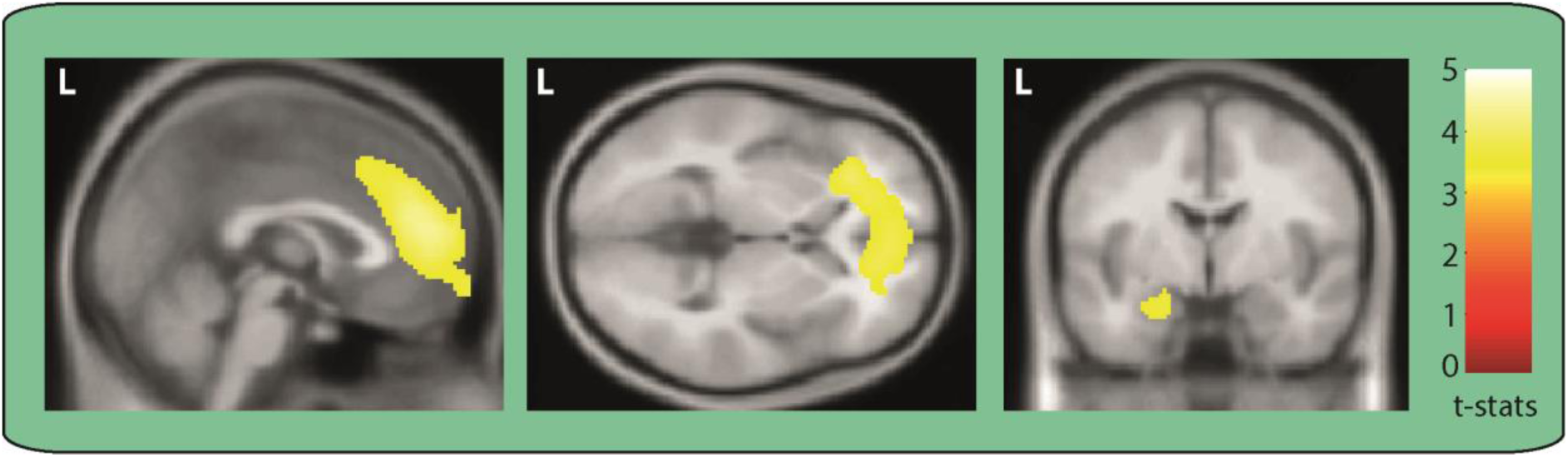
Engagement of vmPFC and left hippocampus during AM retrieval. MEG source reconstruction of broadband (1-30 Hz) power changes during AM retrieval compared to the counting baseline condition. We examined the first 5 secs of each trial when participants would likely have been most fully engaged in recalling their AMs or mentally counting (highly similar results were obtained when the full 10 sec task periods were examined). Images are superimposed on the Montreal Neurological Institute 152 T1 MR image. L=left hemisphere.

### vmPFC Leads and Drives Activity in the Hippocampus During the Initiation of AM Retrieval

Having identified that our two regions of interest exhibited significant power changes during AM retrieval, we next examined what happened during the initiation of AM recall. As outlined above, a key interest was in the temporal order of vmPFC and hippocampal engagement, and whether the vmPFC instigates retrieval. To address this question, we leveraged the high temporal precision of MEG and examined averaged event-related signals for the AM retrieval and baseline counting conditions for both vmPFC and hippocampal channels.

Overlaying the event-related signals of the two channels suggested a strong temporal order effect during AM retrieval but not baseline counting (Figure 3B). During AM retrieval, the maximum response of the vmPFC occurred significantly before the maximum response of the hippocampus. We extracted the maximum response of the vmPFC and hippocampus for each participant and compared their latencies (Figure 3C). The maximum response of the vmPFC occurred on average 125.6 msec (SD=36.2) after AM retrieval onset, while the maximum response of the hippocampus occurred around 65 msec later, at 190.3 msec (SD=41.4; t(df=34)=4.9, p<0.0001). This temporal dissociation was not evident for the baseline counting condition, where the temporal order of vmPFC and hippocampal maximum responses were randomly distributed across participants (vmPFC: Mean=160.2 msec, SD=40.1); HPC: Mean=177.6 msec, SD=41.1; t(df=34)=1.3, p<0.21). These analyses suggest that the vmPFC may initiate AM retrieval. However, while these event-related signals lend support to the idea of a causal relationship, they are not conclusive. Therefore, in a follow-up analysis we examined the effective connectivity between the event-related signals generated by the vmPFC and hippocampus, asking whether one region exerted a directing influence over the other.

**Figure 3.**
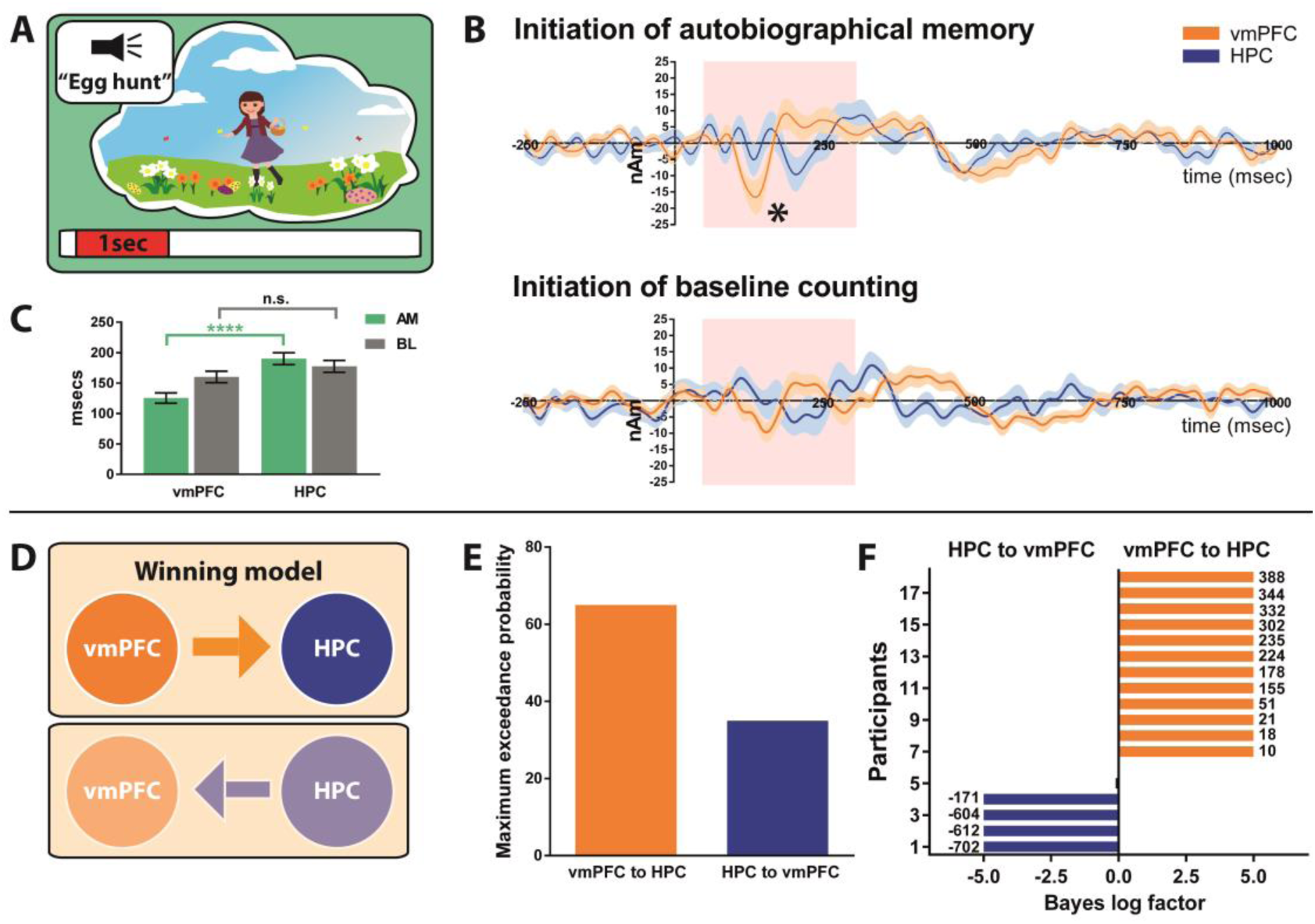
Initiation of AM retrieval. (*A*) Schematic of the <1 sec AM retrieval initiation period under consideration. (*B*) Event-related signals for AM retrieval and baseline counting for vmPFC (in orange) and the left hippocampus (in blue). The continuous lines represent the mean and the shaded areas around the lines represent the SEM. The pink shaded boxes highlight the period from 50 to 300 msec in which the maximum response was examined. *=significant difference between vmPFC and the left hippocampus engagement. (*C*) Bar graph displaying the means and SEM of the maximum responses for AM retrieval (green) and baseline counting (grey) for the vmPFC and the left hippocampus. Whereas during AM retrieval, the maximum response occurred around 100 msec after cue onset, the maximum response of the left hippocampus occurred reliably later; ***=p<0.0001. No such temporal order effect was observed during baseline counting. (*D*) Two proposed models of effective connectivity between the vmPFC and left hippocampus. (*E*) Results of Bayesian model comparison indicated a stronger influence of the vmPFC on left hippocampal activity during AM retrieval initiation. (*F*) Log Bayes factor for each participant. Orange bars indicate positive to strong evidence for vmPFC driving left hippocampal activity, the model which was most consistent across participants. Blue bars represent the four participants where evidence of the left hippocampus driving vmPFC activity was observed. Black bars show the remaining two participants where there was no conclusive evidence for either model. Where the log Bayes factor exceeded five, bars are truncated and exact values are adjacently displayed. AM=Autobiographical memory, BL=baseline counting, vmPFC=ventromedial prefrontal cortex, HPC=hippocampus.

Event-related changes can be viewed as perturbations of cortical networks and explained by underlying changes in effective connectivity or coupling among neural sources. Here we used DCM to elucidate the most likely biological networks underpinning the event-related signals (Garrido et al. 2007). We compared two hypotheses, one where the vmPFC influenced hippocampal activity and another where the hippocampus influenced vmPFC activity (Figure 3D). As Figure 3E illustrates, the model most likely to be the winning model across participants, with a probability of 65.0%, was the vmPFC exerting causal influence over the anterior hippocampal during the initiation of AM retrieval. This was the most consistent model across participants (Figure 3F) as indicated by a log Bayes factor >3.

### vmPFC Drives Activity in the Hippocampus Over the Duration of AM Retrieval

Having examined the initiation of AM recall, we next asked whether the driving influence of the vmPFC over the hippocampus was sustained over the course of retrieval. We examined the first 5 secs of each trial when participants were likely to be most fully engaged in recalling their AMs or mentally counting (of note, highly similar results were obtained when the full 10 sec task periods were examined). To examine effective connectivity in this context, we used DCM for CSD (Kiebel et al. 2009; Moran et al. 2009), a technique that infers model parameters and model evidence based upon cross spectra of MEG data across different regions. This method is especially suited to interrogating broadband signals (as in our case, 1-30 Hz) within which cross spectral densities are often manifest. As before, we specified two models, one where information was allowed to flow from the vmPFC to the hippocampus, and another model where information flowed from the hippocampus to the vmPFC (Figure 4B). The model most likely to be the winning model, with a probability of 92.7%, was the vmPFC exerting a causal influence over the hippocampus during AM retrieval (Figure 4C). This outcome was consistent for the majority of participants, with only 2 participants showing evidence for the model where hippocampus drove vmPFC activity (Figure 4D). Our effective connectivity analysis, therefore, indicated that the vmPFC directed hippocampal activity throughout the AM retrieval process.

**Figure 4.**
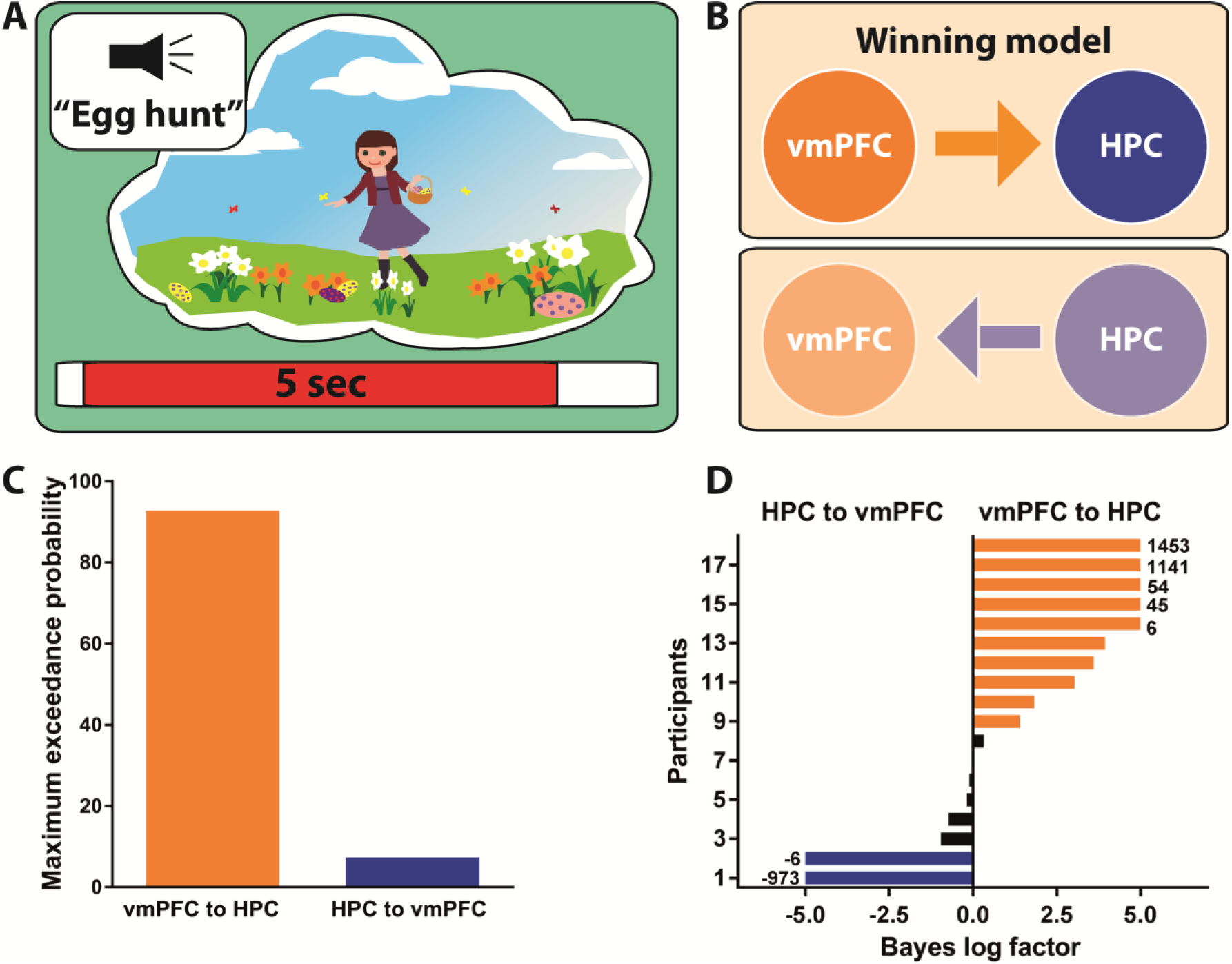
Effective connectivity between vmPFC and left hippocampus during AM retrieval. (*A*) Schematic of the 5 sec AM retrieval period under consideration. (*B*) Two proposed models of effective connectivity between the vmPFC and the left hippocampus. (*C*) Results of Bayesian model comparison indicated that the ventromedial prefrontal cortex (vmPFC) influenced activity in the left hippocampus (HPC) during AM retrieval. (*D*) Log Bayes factor for each participant. Orange bars indicate positive to strong evidence for vmPFC driving left hippocampal activity, the model which was most consistent across participants. Blue bars represent the two participants where evidence of the left hippocampus driving vmPFC activity was observed. Black bars show the remaining participants where there was no conclusive evidence for either model. Where the log Bayes factor exceeded five, bars are truncated and exact values are adjacently displayed.

In summary, honing in on the very earliest initiation phase of AM retrieval using an event related analysis, we found that the vmPFC engaged ∼65 msec earlier than the hippocampus and appeared to exert a causal influence over hippocampal activity at the beginning of each AM trial. Moreover, during a prolonged phase of AM retrieval during which participants were engaged in vivid, detail-rich AM retrieval, the vmPFC also drove activity in the hippocampus over this extended period.

### AMs – the Effect of Remoteness

As alluded to in the Introduction, there are different views about the involvement of the vmPFC and hippocampus in supporting memories as they age (Sekeres et al. 2018a,b; Barry and Maguire 2019a,b; Moscovitch and Nadel, 2019). We addressed this issue by splitting the AMs into different age categories. For each participant, 12 AMs were less than 1 month old (<1M), 12 were between 4 and 12 months old (4-12M), 12 were between 16 and 20 months old (16-20M), and 12 were between 2 and 5 years old (2-5Y) (Barry et al. 2018).

Examining their phenomenological qualities, we found no significant differences between the age categories for vividness (F(df=17)=0.05, p=0.94), ease of recall (F(df=17)=0.26, p=0.71), personal significance (F(df=17)=3.0, p=0.06), and frequency of rehearsal (F(df=17)=0.9, p=0.43). There was a significant effect of emotional valence (F(df=17)=5.88, p=0.002), whereby 2-5Y old memories were rated as more positive than the other AM age categories (**<1M**, t(df=17)=3.3, p=0.02, **4-12M**, t(df=17)=3.1, p=0.04, and **16-20M**, t(df=17)=3.7, p=0.01).

### vmPFC and Hippocampal Neural Dynamics Support AM Retrieval Irrespective of Remoteness

We next determined which brain regions were engaged during AM retrieval by estimating the difference in broadband power (1-30 Hz) between the AM and counting baseline conditions in source space for each AM age category. The most significant power changes were observed in two brain regions – the vmPFC (peak MNI coordinate: **<1M** 6, 60, −10, t=3.30; **4-12M** 14, 60, −12, t=3.33; **16-20M** 10, 60, −12, t=3.19, **2-5Y** 6, 54, −10, t=3.21) and left anterior hippocampus (**<1M** −20, −8, −24, t=3.91; **4-12M** −18, −10, −22, t=3.17, **16-20M** −20, −6, −28, t=3.71, **2-5Y** −20, −4, −26, t=2.95) for all age categories of AM when compared to the baseline counting task (Figure 5). As before, changes in both regions represented an attenuation of power during AM retrieval. Our data, therefore, suggest that both the vmPFC and hippocampus are engaged during AM retrieval irrespective of remoteness.

**Figure 5.**
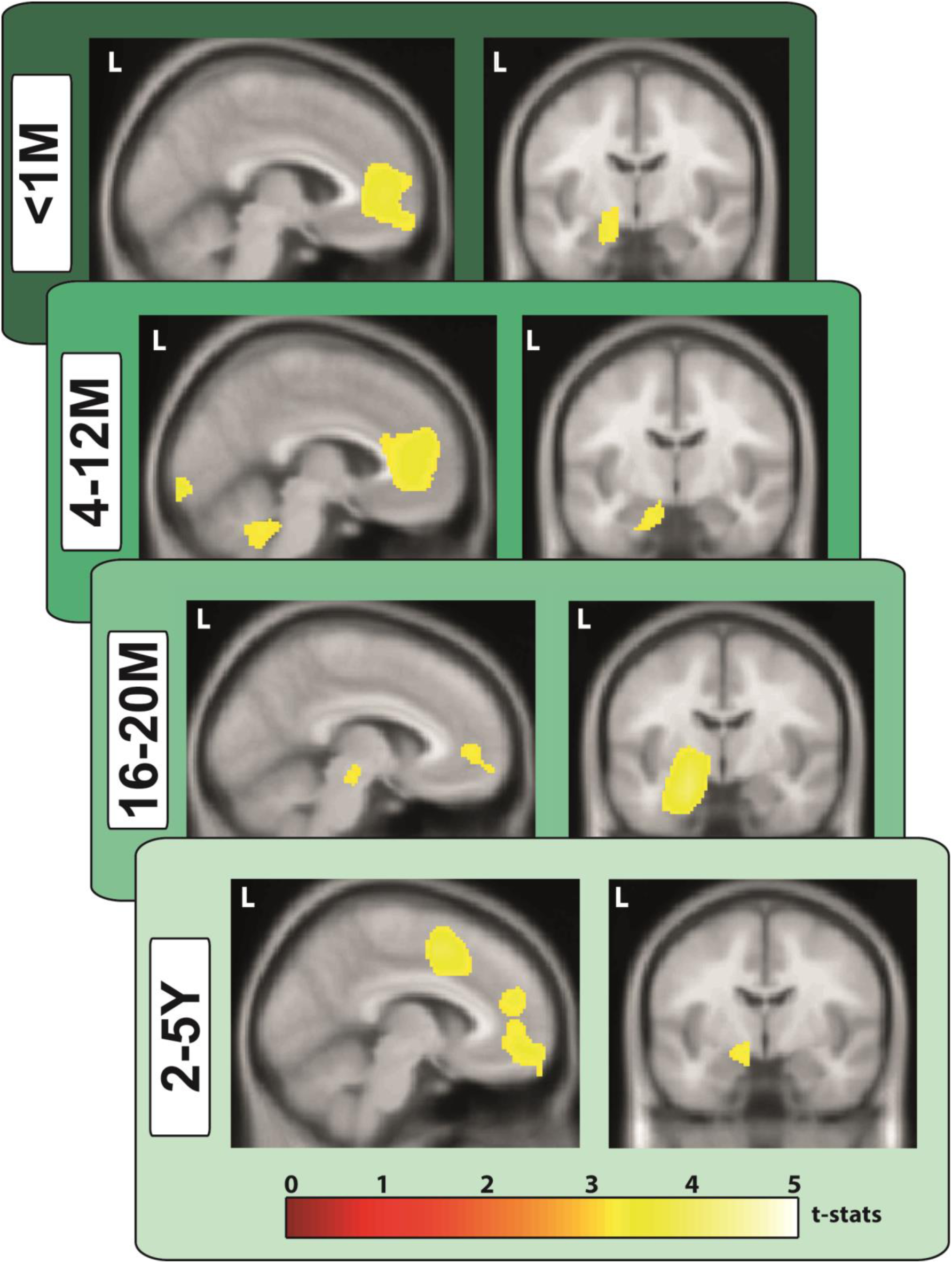
Engagement of vmPFC and left hippocampus during retrieval of AMs of different ages. MEG source reconstruction of broadband (1-30 Hz) power changes during AM retrieval compared to the counting baseline condition. We examined the first 5 secs of each trial when participants were most likely to be fully engaged in recalling their AMs or mentally counting (highly similar results were obtained when the full 10 sec task periods were examined). Images are superimposed on the Montreal Neurological Institute 152 T1 MR image. L=left hemisphere, M=months, Y=years.

### Initiation of AM Retrieval – an Effect of Remoteness

We then examined whether the instigation of AM retrieval was affected by memory remoteness. We generated event-related signals for vmPFC and hippocampal activity during the initiation of retrieval for each age category of AM. Plotting the event-related signal traces for both channels against each other suggested that the maximum response did not significantly differ between vmPFC and hippocampus for very recent memories (**<1M**, vmPFC: Mean=140 msec, SD=33; HPC: Mean=155 msec, SD=25; t(df=17)=1.54, p=0.13; Figure 6A). However, for the other memory age categories, the maximum response of the vmPFC occurred significantly earlier than that of the hippocampus (**4-12M** vmPFC: Mean=135 msec, SD=26; HPC: Mean=195 msec, SD=25; t(df=17)=7.06, p<0.0001; **16-20M** vmPFC: Mean=140 msec, SD=23; HPC: Mean=190 msec, SD=26; t(df=17)=6.10, p<0.0001; **2-5Y** vmPFC: Mean=130 msec, SD=24; HPC: Mean=185 msec, SD=23; t(df=17)=7.02, p<0.0001). Of note, while the vmPFC showed consistent timing of responses for all memory ages (Figure 6B orange bars), the hippocampus showed a comparable early response for the <1M old memories, but then for all other memory ages, the response lagged significantly behind that of the vmPFC (Figure 6B blue bars). This effect was confirmed by statistical analyses with no effect for the latency of the vmPFC response (F(df=17)=0.57, p=0.64), but a significant main effect for the latency of the hippocampal response (F(df=17)=19.65, p<0.0001).

**Figure 6.**
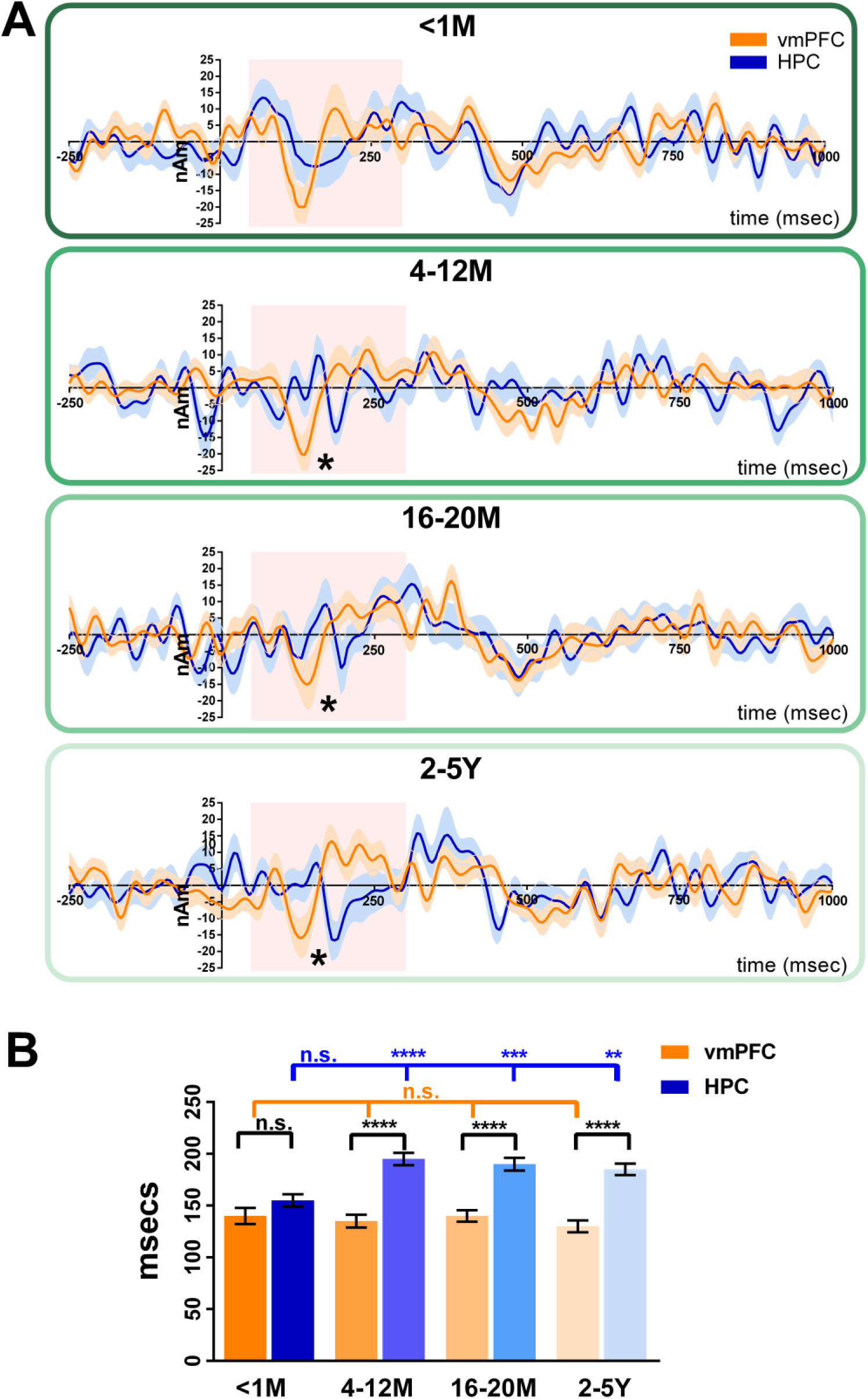
Initiation of retrieval – the effect of AM remoteness. (*A*) Event-related signals for AM retrieval and baseline counting for vmPFC (in orange) and the left hippocampus (in blue). The continuous lines represent the mean and the shaded areas around the lines represent the SEM. The pink shaded boxes highlight the period from 50 to 300 msec in which the maximum response was examined. *=significant difference between vmPFC and left hippocampus engagement (with Bonferroni correction at p<0.01). (*B*) Bar graph displaying the means and SEM of the maximum responses for AM for the vmPFC (orange bars) and the left hippocampus (blue bars). For AMs <1M old, the maximum response of the vmPFC and left hippocampus occurred at around the same time. For all other AM ages, the maximum response of the vmPFC occurred significantly earlier than the left hippocampus. vmPFC=ventromedial prefrontal cortex, HPC=hippocampus, M=months, Y=years, ns=no statistically significant difference, **=p<0.01, ***=p<0.001, ****=p<0.0001.

We then assessed whether the causal relationship between vmPFC and hippocampus during the initiation of AM retrieval was affected by memory remoteness using DCM for event-related signals (Figure 7B). Overall, the model most likely to be the winning model was the vmPFC exerting causal influence over the hippocampus indicated by a significantly higher (i.e., less negative) free energy value than the reverse model (vmPFC to HPC, Mean=-1394, SD=440.1; HPC to vmPFC, Mean=-1642, SD=187.5; t(df=17)=2.28, p=0.036; Figure 7C). Examining the connectivity strengths revealed a significant effect of condition (F(df=17)=5.21, p=0.001), whereby the driving influence of the vmPFC over the hippocampus was generally stronger during AM retrieval than during baseline counting. Furthermore, while connectivity during baseline counting was not different from zero, connectivity strength was increased for all memory ages (**<1M** Mean=0.19, SD=3.5, t(df=17)=2.2, p=0.041; **4-12M** Mean=0.19, SD=2.8, t(df=17)=2.9, p=0.01; **16-20M** Mean=0.26, SD=2.3, t(df=17)=4.5, p=0.0004; **2-5Y** Mean=0.27, SD=2.6, t(df=17)=4.1, p=0.0008; Figure 7D). Of note, there was no significant difference between connectivity strengths across memory ages (F(df=17)=0.36, p=0.71). Therefore, a stronger casual influence of vmPFC over hippocampus was evident for the initiation of AM retrieval for all memory ages, even for the <1M old memories for which the latencies did not differ between vmPFC and hippocampus.

**Figure 7.**
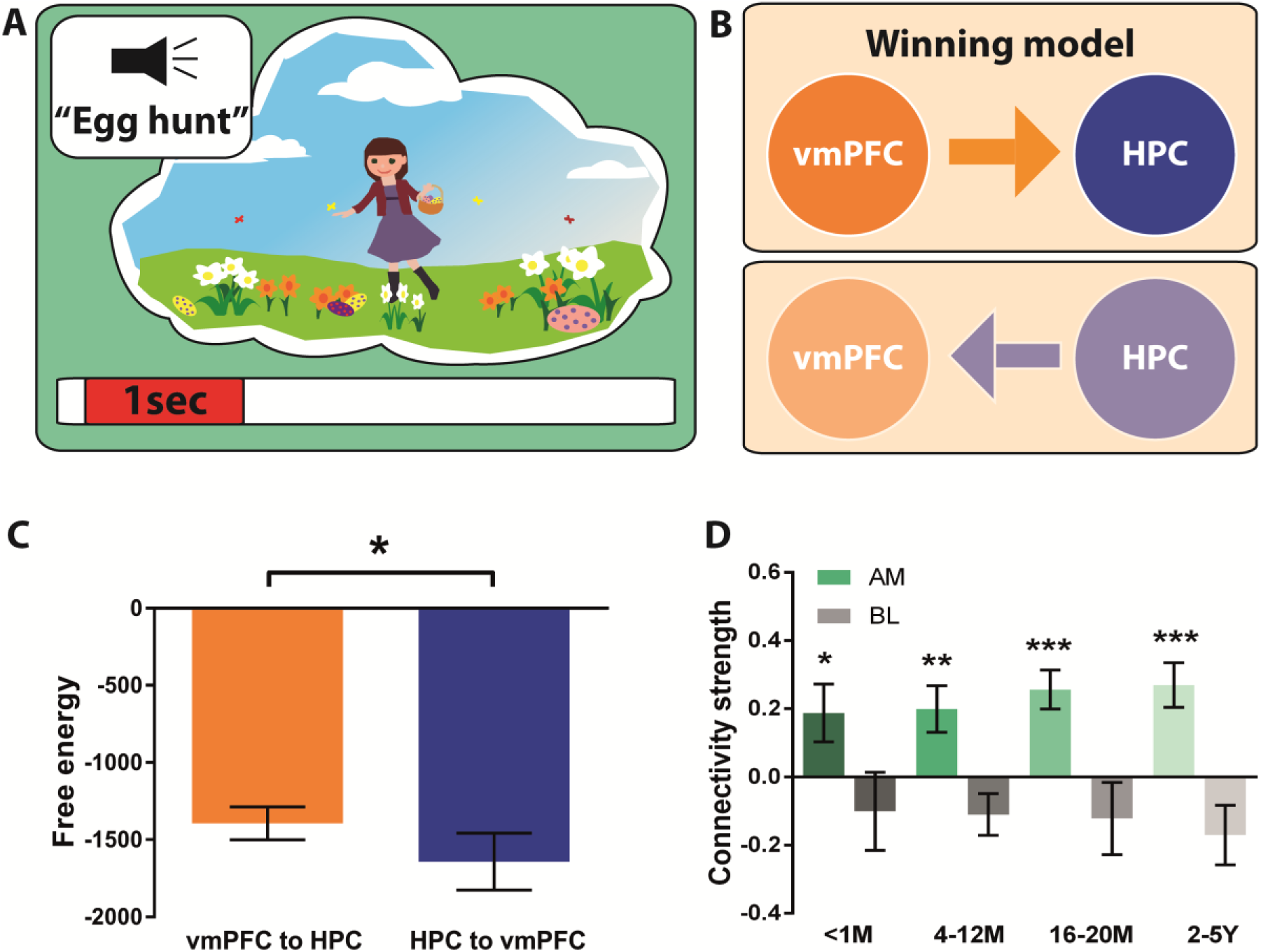
Effective connectivity between vmPFC and left hippocampus during the initiation of retrieval for different AM ages. (*A*) Schematic of the <1 sec AM retrieval initiation period under consideration. (*B*) Two proposed models of effective connectivity between the vmPFC and the left hippocampus. (*C*) Free energy as a measure of model fit indicated a stronger influence of the vmPFC on left hippocampal activity during the initiation of AM retrieval (less negative=more free energy). (*D*) Connectivity strength for all memory ages (green) and the baseline counting condition (grey). Means and SEM are displayed. For all memory ages, the most likely best fitting model was the vmPFC exerting influence over the left hippocampus. AM=Autobiographical memory, BL=baseline counting, vmPFC=ventromedial prefrontal cortex, HPC=hippocampus, M=months, Y=years. *=p<0.05, **=p<0.01, ***=p<0.001.

### vmPFC Drives Activity in the Hippocampus Over the Duration of AM Retrieval Irrespective of Remoteness

We next examined whether the driving influence of the vmPFC over the hippocampus was sustained over the course of retrieval for the different AM age categories using DCM for CSD (Figure 8B). Overall, the model most likely to be the winning model was the vmPFC exerting causal influence over the hippocampus, indicated by a significantly higher (i.e., less negative) free energy value than the reverse model (vmPFC to HPC, Mean=-426.7, SD=669.4; HPC to vmPFC, Mean=-747.8, SD=551.1; t(df=17)=2.21, p=0.042; Figure 8C). Examining the connectivity strengths revealed a significant effect of condition (F(df=17)=3.3, p=0.015), whereby the driving influence of the vmPFC over hippocampal activity was stronger during AM retrieval than during baseline counting. Furthermore, while connectivity during baseline counting was not different from zero, connectivity strength was increased for all memory ages (**<1M** Mean=0.26, SD=3.4, t(df=17)=3.2, p=0.006; **4-12M** Mean=0.32, SD=3.8, t(df=17)=3.4, p=0.004; **16-20M** Mean=0.32, SD=4.2, t(df=17)=3.3, p=0.014; **2-5Y** Mean=0.37, SD=4.3, t(df=17)=3.5, p=0.003; Figure 8D). Of note, there was no significant difference in connectivity strengths across memory ages (F(df=17)=0.2, p=0.87). Therefore, a stronger casual influence of vmPFC over hippocampus for the duration of retrieval was evident for all AMs irrespective of remoteness.

**Figure 8.**
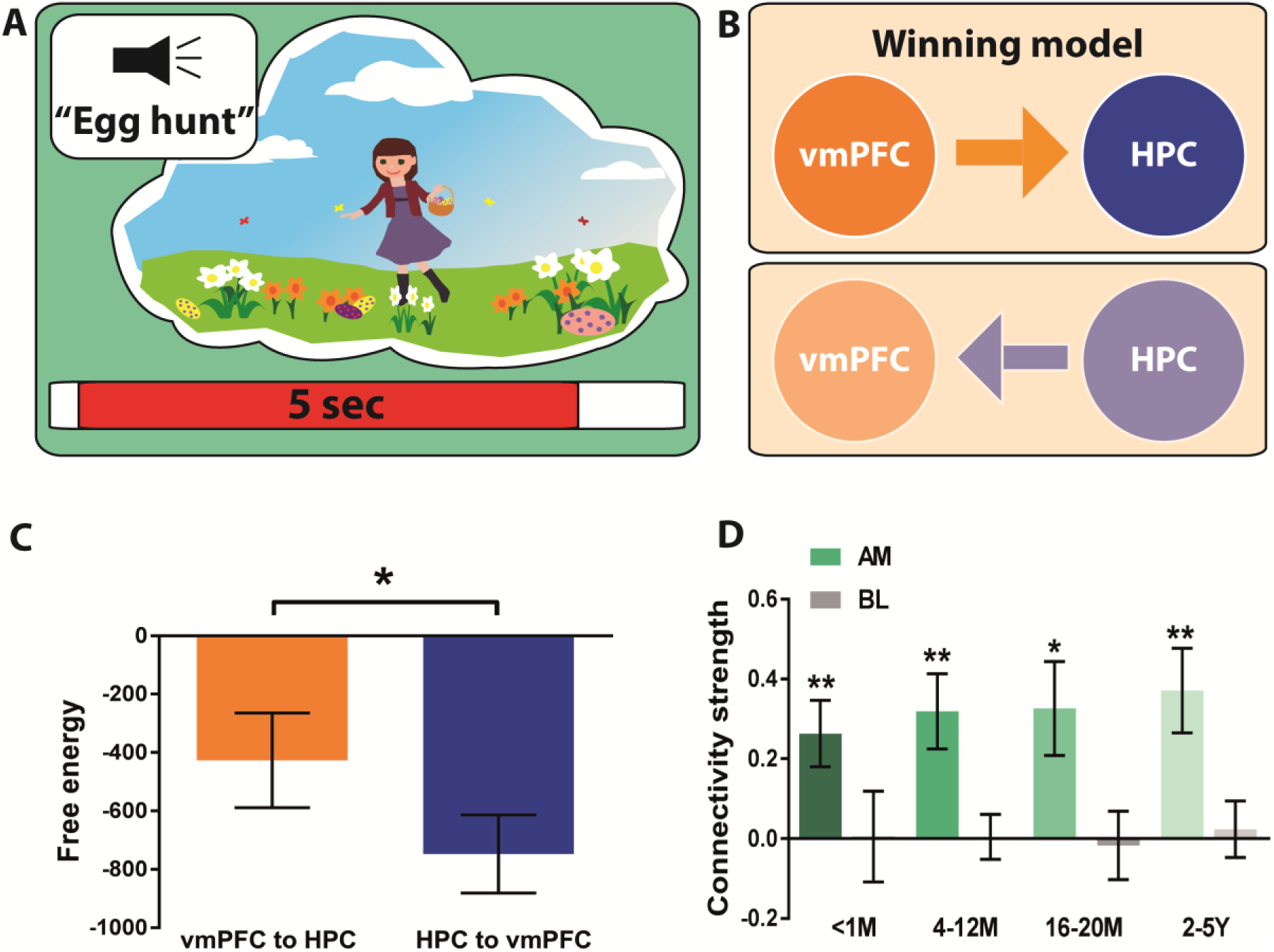
Effective connectivity between vmPFC and left hippocampus during AM retrieval for different memory ages. (*A*) Schematic of the 5 sec AM retrieval period under consideration. *(B)* Two proposed models of effective connectivity between the vmPFC and left hippocampus. *(C)* Free energy as a measure of model fit indicated a stronger influence of the vmPFC on left hippocampal activity during AM retrieval (less negative=more free energy). (*D*) Connectivity strength for all memory ages (green) and the baseline counting condition (grey). Means and SEM are displayed. For all memory ages, the most likely best fitting model was the vmPFC exerting influence over the left hippocampus. AM=Autobiographical memory, BL=baseline counting, vmPFC=ventromedial prefrontal cortex, HPC=hippocampus, M=months, Y=years. *=p<0.05, **=p<0.01.

## Discussion

Autobiographical memories provide the continuity in life’s narrative, are the vehicle for much of our knowledge acquisition, and allow us to live independently. Despite being central to everyday mental life, there is no agreed model of AM retrieval, and there is a lack of understanding about the neural mechanisms involved. In this study we set out to provide missing information that is fundamental for helping to elucidate how neural responses in two key brain regions, the vmPFC and hippocampus, lead to an ability to seamlessly recall the past. Leveraging the high temporal resolution of MEG, we report: (1) electrophysiological evidence that the vmPFC and left hippocampus were engaged during AM retrieval, showing the greatest power changes across the whole brain; (2) that responses in vmPFC preceded activity in the left hippocampus during initiation of AM recall, except during retrieval of the most recent AMs; (3) vmPFC drove left hippocampal activity during recall initiation, and also as AMs unfolded over subsequent seconds; and (4) this hierarchical relationship, with vmPFC driving hippocampus, was evident regardless of AM age. We discuss each of these findings in turn.

### Electrophysiological Evidence of vmPFC and Hippocampal Involvement in AM Retrieval

Only a small number of previous studies have used MEG to examine AM recall. Along with Fuentemilla et al. (2014) and Hebscher et al. (2019), we found that the hippocampus and vmPFC were engaged during AM retrieval. Moreover, we observed that these regions showed the greatest AM-related power changes across the brain relative to a baseline task. Interestingly, AM recollection was associated with attenuation of power. This aligns with accumulating evidence from electroencephalography (Fellner et al. 2017) and MEG (Guderian et al. 2009) demonstrating a strong decrease in medial temporal lobe power during episodic memory encoding. These findings have been validated using direct intracranial recordings in humans, with brain-wide decreases in theta power predicting subsequent recall (Burke et al. 2013; Greenberg et al. 2015), including in the hippocampus (Sederberg et al. 2007; Lega et al. 2012; Matsumoto et al. 2013; Lega et al. 2017). A decrease in power has also been reported during episodic memory retrieval (Michelmann et al. 2016) and during the imagination of novel scenes (Barry et al. 2019a,b).

While several of the above studies focused on theta band oscillations, other reports have documented memory-related effects in the alpha, beta, gamma and even high-gamma bands, using various approaches including phase-frequency coupling, enveloping, and other combinations of interactions between frequencies (Sederberg et al. 2003; Mormann et al. 2005; Sederberg et al. 2007; Axmacher et al. 2008; Axmacher et al. 2010; Fell and Axmacher 2011). Therefore, in the current study, we did not constrain our analyses to a single frequency band but opted instead to examine a broadband signal. This also benefitted the cross-spectral density DCM analysis, because the models therein test interactions across frequency bands. Of note, a beamformer source analysis constrained to the theta band resulted in highly similar findings.

### vmPFC Leads and Drives Hippocampal Activity During AM Recall

Previous DCM fMRI studies have shown that during AM recall elaboration, the vmPFC drove the hippocampus when generic cues were used to trigger recall (St Jacques et al. 2011; Nawa and Ando 2019). However, fMRI has relatively poor temporal resolution, thus prohibiting interpretations of the precise temporal order of neural events. In their MEG study, Fuentemilla et al. (2014) noted phase coupling between hippocampus and vmPFC, but no MEG study has compared the exact timings of hippocampus and vmPFC engagement, nor the effective connectivity between them, despite the important implications for a mechanistic understanding of AM. The current study, therefore, represents the first attempt to decipher the precise temporal order of neural events during AM retrieval at the source level while using highly specific memory cues.

We found that the vmPFC engaged significantly earlier than the left hippocampus, with this effect emerging during AM retrieval but not during a baseline task. It was most apparent between 120-200msec after cue onset which, at first glance, seems very early. However, it is pertinent to bear in mind our experimental design when considering the time course of the neural responses. Participants were seated in the MEG scanner and on each trial had already received the instruction to close their eyes just before a memory cue was provided. Hence, they were in a state of readiness to recall. Moreover, the participants themselves had generated the two-word memory cues, a unique title for each memory. Therefore, once the unique auditory memory cue started, and while it was still playing, they could already begin memory retrieval. This was not a process that waited until cue offset. We are not arguing that by 120ms participants were recalling fully elaborated autobiographical memories. This time window captured the initiation of the retrieval process, presumably involving the identification of the appropriate memory and instigation of its recall.

Early frontal lobe responses are not without precedent. Using intracranial EEG, frontal responses have been reported to appear almost instantaneously, with the hippocampus lagging significantly behind (Sederberg et al. 2003; Sederberg et al. 2007). Moreover, a recent MEG study examining the temporal dynamics during AM retrieval in sensor space reported similarly early event-related potentials over frontal sensors at around 100-200msec (Hebscher et al. 2020). The authors of this latter study proposed that these early frontal responses may reflect the structural analysis of the stimuli. While this could be the case, we suggest that this early response may also signify the vmPFC starting to drive downstream processes, including those in the hippocampus.

Related to this point, when we examined the direction of information flow between hippocampus and vmPFC at this very early initiation stage of AM recall, we found that the vmPFC drove hippocampal event-related signals. This influence of vmPFC over hippocampus echoes that documented previously in relation to the imagination of novel scenes. Barry et al. (2019a) found that the vmPFC engaged earlier and drove hippocampal activity during the creation of novel scene imagery. Similarly, aligning with previous fMRI reports (St Jacques et al. 20110; Nawa and Ando 2019), we found that the vmPFC also exerted directional influence over the hippocampus in the seconds following AM recall initiation, when participants were maximally engaged in elaborating their personal memories. Of note, the memory cues in our study were highly specific, and so our findings cannot be explained by the vmPFC leading retrieval merely because cues were generic and lacked specificity, as proposed by one account of memory recall (Robin and Moscovitch 2017). Moreover, the engagement of posterior cortical areas predicted by the same account during the elaboration phase was not evident in our data. Given the vivid and detailed nature of the autobiographical memories included in our study, and the high success with which participants vividly recalled them (indexed by the ratings made for each trial during scanning), this is unlikely to be explained by weak elaboration. Instead our data reveal that the interplay between vmPFC and hippocampus is central to AM retrieval, with the vmPFC directing this interaction from the start and throughout.

### vmPFC Leads and Drives Hippocampal Activity Regardless of AM remoteness

While cellular studies indicate that the hippocampus does not seem to store anything in the longer term (reviewed in Barry and Maguire 2019a,b), there is an abundance of evidence showing that it nevertheless supports the retrieval of vivid, detail-rich AMs regardless of their age (Gilboa et al. 2004; Bonnici et al. 2012; Sheldon and Levine 2013; Clark and Maguire 2016; Bonnici and Maguire 2018; McCormick et al. 2018b; Sekeres et al. 2018b; Miller et al. 2020). Recent and remote memories are also represented in the vmPFC (Bonnici et al. 2012), with several longitudinal fMRI studies indicating that their detectability increases with remoteness (Bonnici et al. 2012; Barry et al. 2018; Bonnici and Maguire 2018). We found that the hippocampus and vmPFC showed the greatest power changes across the whole brain during retrieval of AMs of any age, ranging from those that were less than one month old to memories that were five years old.

One difference that did emerge concerned the temporal order of vmPFC and hippocampal engagement during very early AM recall initiation. For the most recent AMs, there was no difference between the two brain regions, however, for all other AMs, a significant timing lag was evident, with the hippocampus slower to respond. Of note, the timing of vmPFC engagement was consistent irrespective of AM age, rather it was the hippocampus that became slower to respond once AMs were no longer very recent. This distinction between recent and more remote AMs cannot be explained by differences in vividness or ease of recall, as these factors did not differ as a function of AM age. Moreover, it is unlikely the effect is cue-related, or can be explained by participants recalling the pre-scan memory-harvesting interview, as these were similar and pertained to all memories irrespective of age. Instead, we suggest these findings accord with the view that recent AMs may still be available in the hippocampus, and so it does not need to await direction from the vmPFC in order to reconstruct them (McCormick et al. 2018a; Ciaramelli et al. 2019; Barry and Maguire 2019a,b).

Our effective connectivity findings also revealed another dimension to hippocampal-vmPFC interactions. Irrespective of AM age, during the very earliest recall initiation period, and over the subsequent seconds as memory events unfolded, the vmPFC exerted a driving influence on hippocampal activity. Of note, even when AMs were recent, and there was no difference in the timing of hippocampal and vmPFC engagement, vmPFC nevertheless still exerted an influence over hippocampal activity. These results suggest that the vmPFC is actively involved in AM processing in the first few weeks after a memory has been formed.

### Conclusions and Theoretical Considerations

The hippocampus and vmPFC are crucial for vivid AM retrieval, however, the precise dynamic interplay between them has remained elusive. Whereas traditional views assume that the hippocampus initiates event recall (Teyler and DiScenna 1986; Teyler and Rudy 2007), an alternative perspective proposes that the vmPFC instigates and coordinates hippocampal-dependent processes (McCormick et al. 2018a; Barry and Maguire 2019a,b). Our findings of vmPFC engaging significantly in advance of hippocampus, and driving oscillatory activity in hippocampus both at the start and throughout memory retrieval, aligns strongly with this latter view. Moreover, that the vmPFC influenced hippocampal activity even during retrieval of recent AMs, provides further insights into systems-level consolidation. The vmPFC may work with the hippocampus early in the consolidation process (Bonnici et al. 2012; Kitamura et al. 2017) to start integrating AMs with existing schema. For AMs that are already consolidated, the vmPFC might draw upon relevant schema to orchestrate AM recall, influencing what information the hippocampus receives and uses to re-construct a past event (McCormick et al. 2018a). Our study was not designed to examine schema, but motivates further MEG research in this domain. In a similar vein, it would also be interesting in future MEG studies to vary other features of autobiographical memories, such as their subjective vividness and the detail with which they are recalled.

Another active area of debate, recently reinvigorated by several opinion pieces (Barry and Maguire, 2019a,b; Moscovitch and Nadel, 2019) concerns whether the hippocampus stores traces of autobiographical memories in the longer-term. While this is undoubtedly an important question, it was not the focus of the current study. Our finding that vmPFC engagement precedes and drives that of the hippocampus does not permit conclusions to be drawn about whether vmPFC activates traces that are present in the hippocampus or initiates reconstruction of a memory anew in the absence of a hippocampal trace. Adjudicating between these options will be challenging and requires a different experimental approach.

In summary, the results of this study re-cast the positions of the hippocampus and vmPFC in the AM retrieval hierarchy by providing new information about their neural responses and how they might give rise to our ability to re-experience past autobiographical events.

## NOTES

We thank David Bradbury for technical assistance and Vladimir Litvak for analysis advice.

*Conflict of Interest:* The authors declare no conflicts of interests.

## AUTHOR CONTRIBUTIONS

C.M. and E.A.M. designed the study with input from G.R.B. C.M. collected the data with assistance from D.N.B. C.M. analysed the data with input from the other authors. C.M. and E.A.M wrote the paper with input from the other authors.

## REFERENCES

Addis DR, McIntosh AR, Moscovitch M, Crawley AP, McAndrews MP. 2004. Characterizing Spatial and Temporal Features of Autobiographical Memory Retrieval Networks: A Partial Least Squares Approach. Neuroimage. 23:1460–1471.

Addis DR, Moscovitch M, McAndrews MP. 2007. Consequences of hippocampal damage across the autobiographical memory network in left temporal lobe epilepsy. Brain. 130:2327–2342.

Axmacher N, Henseler MM, Jensen O, Weinreich I, Elger CE, Fell J. 2010. Cross-frequency coupling supports multi-item working memory in the human hippocampus. Proc Natl Acad Sci USA. 107:3228–3233.

Axmacher N, Schmitz DP, Wagner T, Elger CE, Fell J. 2008. Interactions between medial temporal lobe, prefrontal cortex, and inferior temporal regions during visual working memory: a combined intracranial EEG and functional magnetic resonance imaging study. J Neurosci. 28:7304–7312.

Backus AR, Schoffelen JM, Szebenyi S, Hanslmayr S, Doeller CF. 2016. Hippocampal-prefrontal theta oscillations support memory integration. Curr Biol. 26:450–457.

Barry DN, Chadwick MJ, Maguire EA. 2018. Nonmonotonic recruitment of ventromedial prefrontal cortex during remote memory recall. PLoS Biol. 16:e2005479.

Barry DN, Barnes GR, Clark IA, Maguire EA. 2019a. The neural dynamics of novel scene imagery. J Neurosci. 39:4375–4386.

Barry DN, Tierney TM, Holmes N, Boto E, Roberts G, Leggett J, Bowtell R, Brookes MJ, Barnes GR, Maguire EA. 2019b. Imaging the human hippocampus with optically-pumped magnetoencephalography. Neuroimage. 203:116192.

Barry DN, Maguire EA. 2019a. Remote memory and the hippocampus: A constructive critique. Trends Cogn Sci. 23:128–142.

Barry DN, Maguire EA. 2019b. Consolidating the case for transient hippocampal memory traces. Trends Cogn Sci. 23:635–636.

Bertossi E, Ciaramelli E. 2016. Ventromedial prefrontal damage reduces mind-wandering and biases its temporal focus. Soc Cog Affect Neurosci. 11:1783–1791.

Bertossi E, Tesini C, Cappelli A, Ciaramelli E. 2016. Ventromedial prefrontal damage causes a pervasive impairment of episodic memory and future thinking. Neuropsychologia. 90:12–24.

Bonnici HM, Chadwick MJ, Lutti A, Hassabis D, Weiskopf N, Maguire EA. 2012. Detecting representations of recent and remote autobiographical memories in vmPFC and hippocampus. J Neurosci. 32:16982–16991.

Bonnici HM, Maguire EA. 2018. Two years later - Revisiting autobiographical memory representations in vmPFC and hippocampus. Neuropsychologia. 110:159–169.

Burke JF, Zaghloul KA, Jacobs J, Williams RB, Sperling MR, Sharan AD, Kahana MJ. 2013. Synchronous and asynchronous theta and gamma activity during episodic memory formation. J Neurosci. 33:292–304.

Catani M, Dell’acqua F, Thiebaut de Schotten M. 2013. A revised limbic system model for memory, emotion and behaviour. Neurosci Biobehav Rev. 37:1724–1737.

Catani M, Dell’acqua F, Vergani F, Malik F, Hodge H, Roy P, Valabregue R, Thiebaut de Schotten M. 2012. Short frontal lobe connections of the human brain. Cortex. 48:273–291.

Ciaramelli E, De Luca F, Monk AM, McCormick C, Maguire EA. 2019. What “wins” in VMPFC: Scenes, situations, or schema? Neurosci Biobehav Rev. 100:208–210.

Ciaramelli E, Ghetti S, Frattarelli M, Ladavas E. 2006. When true memory availability promotes false memory: evidence from confabulating patients. Neuropsychologia. 44:1866–1877.

Clark IA, Maguire EA. 2020. Do questionnaires reflect their purported cognitive functions? Cognition 195:104114.

Clark IA, Maguire EA. 2016. Remembering preservation in hippocampal amnesia. Annu Rev Psychol. 67:51–82.

Dalal SS, Jerbi K, Bertrand O, Adam C, Ducorps A, Schwartz D, Garnero L, Baillet S, Martinerie J, Lachaux J-P. 2013. Evidence for MEG detection of hippocampus oscillations and cortical gamma-band activity from simultaneous intracranial EEG. Epilep Behav 28: 310–311.

Della Sala S, Laiacona M, Spinnler H, Trivelli C. 1993. Autobiographical recollection and frontal damage. Neuropsychologia. 31:823–839.

Fell J, Axmacher N. 2011. The role of phase synchronization in memory processes. Nat Rev Neurosci. 12:105–118.

Felleman DJ, Van Essen DC. 1991. Distributed hierarchical processing in the primate cerebral cortex. Cereb Cortex. 1:1–47.

Fellner MC, Volberg G, Wimber M, Goldhacker M, Greenlee MW, Hanslmayr S. 2017. Spatial mnemonic encoding: Theta power decreases and medial temporal lobe BOLD increases co-occur during the usage of the method of loci. eNeuro. 3:ENEURO.0184-0116.2016.

Friston KJ, Harrison L, Penny W. 2003. Dynamic causal modelling. Neuroimage. 19:1273–1302.

Fuentemilla L, Barnes GR, Duzel E, Levine B. 2014. Theta oscillations orchestrate medial temporal lobe and neocortex in remembering autobiographical memories. Neuroimage. 85:730–737.

Fuentemilla L, Palombo DJ, Levine B. 2018. Gamma phase-synchrony in autobiographical memory: Evidence from magnetoencephalography and severely deficient autobiographical memory. Neuropsychologia. 110:7–13.

Garrido MI, Kilner JM, Kiebel SJ, Stephan KE, Friston KJ. 2007. Dynamic causal modelling of evoked potentials: a reproducibility study. Neuroimage. 36:571–580.

Gilboa A, Marlatte H. 2017. Neurobiology of schemas and schema-mediated memory. Trends Cogn Sci. 21:618–631.

Gilboa A, Winocur G, Grady CL, Hevenor SJ, Moscovitch M. 2004. Remembering our past: functional neuroanatomy of recollection of recent and very remote personal events. Cereb Cortex. 14:1214–1225.

Greenberg JA, Burke JF, Haque R, Kahana MJ, Zaghloul KA. 2015. Decreases in Theta and Increases in High Frequency Activity Underlie Associative Memory Encoding. Neuroimage. 114:257–263.

Guderian S, Schott BH, Richardson-Klavehn A, Duzel E. 2009. Medial temporal theta state before an event predicts episodic encoding success in humans. Proc Natl Acad Sci USA. 106:5365–5370.

Hebscher M, Ibrahim C, Gilboa A. 2020. Precuneus stimulation alters the neural dynamics of autobiographical memory retrieval. Neuroimage. 210:116575.

Hebscher M, Meltzer JA, Gilboa A. 2019. A causal role for the precuneus in network-wide theta and gamma oscillatory activity during complex memory retrieval. eLife. 8:e43114.

Inman CS, James GA, Vytal K, Hamann S. 2018. Dynamic changes in large-scale functional network organization during autobiographical memory retrieval. Neuropsychologia. 110:208–224.

Kaplan R, Doeller CF, Barnes GR, Litvak V, Duzel E, Bandettini PA, Burgess N. 2012. Movement-related theta rhythm in humans: coordinating self-directed hippocampal learning. PLoS Biol. 10:e1001267.

Kiebel SJ, Garrido MI, Moran R, Chen CC, Friston KJ. 2009. Dynamic causal modeling for EEG and MEG. Hum Brain Mapp. 30:1866–1876.

Kitamura T, Ogawa SK, Roy DS, Okuyama T, Morrissey MD, Smith LM, Redondo RL, Tonegawa S. 2017. Engrams and circuits crucial for systems consolidation of a memory. Science. 356:73–78.

Kleist K, Leonhard K, Schwab H. 1940. Die Katatonie auf Grund katamnestischer Untersuchungen. Zeitschrift fuer die gesamte Neurologie und Psychiatrie. 168:535–586.

Lega BC, Germi J, Rugg M. 2017. Modulation of oscillatory power and connectivity in the human posterior cingulate cortex supports the encoding and retrieval of episodic memories. J Cogn Neurosci. 29:1415–1432.

Lega BC, Jacobs J, Kahana M. 2012. Human hippocampal theta oscillations and the formation of episodic memories. Hippocampus. 22:748–761.

Maguire EA, Mullally SL. 2013. The hippocampus: A manifesto for change. J Exp Psychol Gen. 142:1180–1189.

Maguire EA. 2001. Neuroimaging studies of autobiographical event memory. Phil Trans Royal Soc, Lond: Biol Sci. 356:1441–1451.

Matsumoto JY, Stead M, Kucewicz MT, Matsumoto AJ, Peters PA, Brinkmann BH, Danstrom JC, Goerss SJ, Marsh WR, Meyer FB, Worrell GA. 2013. Network oscillations modulate interictal epileptiform spike rate during human memory. Brain. 136:2444–2456.

McCormick C, Ciaramelli E, De Luca F, Maguire EA. 2018a. Comparing and contrasting the cognitive effects of hippocampal and ventromedial prefrontal cortex damage: A review of human lesion studies. Neuroscience. 15:295–318.

McCormick C, Moscovitch M, Valiante TA, Cohn M, McAndrews MP. 2018b. Different neural routes to autobiographical memory recall in healthy people and individuals with left medial temporal lobe epilepsy. Neuropsychologia. 110:26–36.

McCormick C, St-Laurent M, Ty A, Valiante TA, McAndrews MP. 2015. Functional and effective hippocampal-neocortical connectivity during construction and elaboration of autobiographical memory retrieval. Cereb Cortex. 25:1297–1305.

McDermott KB, Szpunar KK, Christ SE. 2009. Laboratory-based and autobiographical retrieval tasks differ substantially in their neural substrates. Neuropsychologia. 47:2290–2298.

Meyer SS, Rossiter H, Brookes MJ, Woolrich MW, Bestmann S, Barnes GR. 2017. Using generative models to make probabilistic statements about hippocampal engagement in MEG. Neuroimage 149:468–482.

Michelmann S, Bowman H, Hanslmayr S. 2016. The temporal signature of memories: Identification of a general mechanism for dynamic memory replay in Humans. PLoS Biol. 14:e1002528.

Miller TD, Chong TT, Aimola Davies AM, Johnson MR, Irani SR, Husain M, Ng TW, Jacob S, Maddison P, Kennard C, Gowland PA, Rosenthal CR. 2020. Human hippocampal CA3 damage disrupts both recent and remote episodic memories. eLife. 24:e41836.

Moran R, Pinotsis DA, Friston K. 2013. Neural masses and fields in dynamic causal modeling. Front Comput Neurosci. 7:1–12.

Moran RJ, Stephan KE, Seidenbecher T, Pape HC, Dolan RJ, Friston KJ. 2009. Dynamic causal models of steady-state responses. Neuroimage. 44:796–811.

Mormann F, Fell J, Axmacher N, Weber B, Lehnertz K, Elger CE, Fernandez G. 2005. Phase/amplitude reset and theta-gamma interaction in the human medial temporal lobe during a continuous word recognition memory task. Hippocampus. 15:890–900.

Moscovitch M. 1992. Memory and working with memory: A component process model based on modules and central systems. J Cogn Neurosci. 4: 257–267.

Moscovitch M, Melo B. 1997. Strategic retrieval and the frontal lobes: evidence from confabulation and amnesia. Neuropsychologia. 35:1017–1034.

Moscovitch M, Nadel L. 2019. Sculpting remote memory: Enduring hippocampal traces and vmPFC reconstructive processes. Trends Cogn Sci. 23:634–635.

Nadel L, Moscovitch M. 1997. Memory consolidation, retrograde amnesia and the hippocampal complex. Curr Opin Neurobiol. 7:217–227.

Nawa NE, Ando H. 2019. Effective connectivity within the ventromedial prefrontal cortex-hippocampus-amygdala network during the elaboration of emotional autobiographical memories. Neuroimage. 189:316–328.

Nolte G. 2003. The magnetic lead field theorem in the quasi-static approximation and its use for magnetoencephalography forward calculation in realistic volume conductors. Phys Med Biol. 48:3637–3652.

Poch C, Fuentemilla L, Barnes GR, Duzel E. 2011. Hippocampal theta-phase modulation of replay correlates with configural-relational short-term memory performance. Journal of Neuroscience. 31:7038–7042.

Robin J, Moscovitch M. 2017. Details, gist and schema: hippocampal–neocortical interactions underlying recent and remote episodic and spatial memory. Curr Opin Behav Sci. 17:114–123.

Robin J, Hirshhorn M, Rosenbaum RS, Winocur G, Moscovitch M, Grady CL. 2015. Functional connectivity of hippocampal and prefrontal networks during episodic and spatial memory based on real-world environments. Hippocampus. 25:81–93.

Scoville WB, Milner B. 1957. Loss of recent memory after bilateral hippocampal lesions. J Neurol Neurosurg Psychiatry. 20:11–21.

Sederberg PB, Kahana MJ, Howard MW, Donner EJ, Madsen JR. 2003. Theta and gamma oscillations during encoding predict subsequent recall. J Neurosci. 23:10809–10814.

Sederberg PB, Schulze-Bonhage A, Madsen JR, Bromfield EB, McCarthy DC, Brandt A, Tully MS, Kahana MJ. 2007. Hippocampal and neocortical gamma oscillations predict memory formation in humans. Cereb Cortex. 17:1190–1196.

Sekeres MJ, Winocur G, Moscovitch M. 2018a. The hippocampus and related neocortical structures in memory transformation. Neurosci Lett. 680:39–53.

Sekeres MJ, Winocur G, Moscovitch M, Anderson JAE, Pishdadian S, Martin Wojtowicz J, St-Laurent M, McAndrews MP, Grady CL. 2018b. Changes in patterns of neural activity underlie a time-dependent transformation of memory in rats and humans. Hippocampus. 28:745–764.

Shallice T, Burgess P. 1996. The domain of supervisory processes and temporal organisation of behaviour. Philos Trans R Soc Lond B Biol Sci. 351:1405–1411.

Sheldon S, Levine B. 2018. The medial temporal lobe functional connectivity patterns associated with forming different mental representations. Hippocampus. 28:269–280.

Sheldon S, Levine B. 2013. Same as it ever was: Vividness modulates the similarities and differences between the neural networks that support retrieving remote and recent autobiographical memories. Neuroimage. 83:880–891.

Spreng RN, Mar RA, Kim AS. 2009. The common neural basis of autobiographical memory, prospection, navigation, theory of mind, and the default mode: a quantitative meta-analysis. J Cogn Neurosci. 21:489–510.

Squire LR. 1992. Memory and the hippocampus: A synthesis from findings with rats, monkeys, and humans. Psych Rev. 99:195–231.

Staudigl T, Hanslmayr S. 2013. Theta oscillations at encoding mediate the context-dependent nature of human episodic memory. Curr Biol. 23:1101–1106.

St Jacques PL, Kragel PA, Rubin DC. 2011. Dynamic neural networks supporting memory retrieval. Neuroimage. 57:608–616.

St-Laurent M, Moscovitch M, Jadd R, McAndrews MP. 2014. The perceptual richness of complex memory episodes is compromised by medial temporal lobe damage. Hippocampus. 24:560–576.

St-Laurent M, Moscovitch M, Tau M, McAndrews MP. 2011. The temporal unraveling of autobiographical memory narratives in patients with temporal lobe epilepsy or excisions. Hippocampus. 21:409–421.

Stephan KE, Penny WD, Daunizeau J, Moran RJ, Friston KJ. 2009. Bayesian model selection for group studies. Neuroimage. 46:1004–1017.

Svoboda E, McKinnon MC, Levine B. 2006. The functional neuroanatomy of autobiographical memory: a meta-analysis. Neuropsychologia. 44:2189–2208.

Teyler TJ, DiScenna P. 1986. The hippocampal memory indexing theory. Behav Neurosci. 100:147–154.

Teyler TJ, Rudy JW. 2007. The hippocampal indexing theory and episodic memory: updating the index. Hippocampus. 17:1158–1169.

Viskontas IV, McAndrews MP, Moscovitch M. 2000. Remote episodic memory deficits in patients with unilateral temporal lobe epilepsy and excisions. J Neurosci. 20:5853–5857.

Williams AN, Ridgeway S, Postans M, Graham KS, Lawrence AD, Hodgetts CJ. 2020. The role of the pre-commissural fornix in episodic autobiographical memory and simulation. Neuropsychologia. doi: 10.1016/j.neuropsychologia.2020.107457

Zeidman P, Maguire EA. 2016. Anterior hippocampus: the anatomy of perception, imagination and episodic memory. Nat Rev Neurosci. 17:173–182.

